# Nanoparticle-based targeted delivery of pentagalloyl glucose reverses elastase-induced abdominal aortic aneurysm and restores aorta to the healthy state in mice

**DOI:** 10.1101/2019.12.16.878017

**Authors:** Saphala Dhital, Naren R. Vyavahare

**Author notes:** **Corresponding Author:** Naren R. Vyavahare, Professor and Hunter Endowed Chair, Department of Bioengineering, 501 Rhodes Research Center, Clemson University, SC, 29634.

## Abstract

**Aim:** Abdominal aortic aneurysms (AAA) is a life-threatening weakening and expansion of the abdominal aorta due to inflammatory cell infiltration and gradual degeneration of extracellular matrix (ECM). There are no pharmacological therapies to treat AAA. We tested the hypothesis that nanoparticle (NP) therapy that targets degraded elastin and delivers anti-inflammatory, anti-oxidative, and ECM stabilizing agent, pentagalloyl glucose (PGG) will reverse advance stage aneurysm in an elastase-induced mouse model of AAA.

**Method and Results:** Porcine pancreatic elastase (PPE) was applied periadventitially to the infrarenal aorta in mice and AAA was allowed to develop for 14 days. Nanoparticles loaded with PGG (EL-PGG-NPs) were then delivered via IV route at 14-day and 21-day (10 mg/kg of body weight). A control group of mice received no therapy. The targeting of NPs to the AAA site was confirmed with fluorescent dye marked NPs and gold NPs. Animals were sacrificed at 28-d. We found that targeted PGG therapy reversed the AAA by decreasing matrix metalloproteinases MMP-9 and MMP-2, and the infiltration of macrophages in the medial layer. The increase in diameter of the aorta was reversed to healthy controls. Moreover, PGG treatment restored degraded elastic lamina and increased the circumferential strain of aneurysmal aorta to the healthy levels.

**Conclusion:** Our results support that site-specific delivery of PGG with targeted nanoparticles can be used to treat already developed AAA. Such therapy can reverse inflammatory markers and restore arterial homeostasis.

## Introduction

An abdominal aortic aneurysm (AAA) is the 13^th^ leading cause of death in the elderly. The common characteristic of AAA disease includes the degradation of the aortic extracellular matrix, smooth muscle cell apoptosis, and gradual weakening and dilation of the aorta^1^. AAA is diagnosed when the aortic diameter is expanded by 50% or more or exceeds 3 cm. In clinical practice, if the diameter reaches 5 cm or more, patients are recommended for surgical intervention. The contributing factors for AAA include male sex, age, genetic factors, hypertension, and smoking history^1 2^.

In AAA, ECM degradation occurs because of the inflammatory process. As the inflammation progresses, activated cells secret pro-matrix metalloproteinases (MMPs). The enzymatic activity of MMPs such as MMP-2, MMP-9, and MMP-12 degrade ECM specifically elastic laminae in the medial layer. Since elastin degradation is one of the first steps during the onset of the AAA, we have been working on developing a drug delivery system that targets degraded elastin at the site of AAA disease. Previously, we have shown that such targeted delivery can deliver agents to reverse calcification of arteries and reverse aortic aneurysms in calcium-chloride (CaCl_2_) injury rat model^3^. We have shown that polyphenols such as pentagalloyl glucose (PGG) and Epigallocatechin gallate (EGCG) can increase elastin deposition by smooth muscle cells derived from healthy or aneurysmal rat aorta^4^. Others have shown than in an elastase model of AAA, a high dose of grape seed polyphenol used orally has a protective role for elastin, and decrease immune cells and MMPs at the AAA site^5^. Green tea polyphenol EGCG was used orally to a rat model of abdominal aortic aneurysm induced by intraluminal infusion of elastase and adventitial simultaneous CaCl_2_ application where EGCG prevented the progression of AAA^5^. These studies used excessively high oral doses of polyphenols at the onset of AAA induction and showed an only protective effect. Moreover, grape seed extracts can have mixtures of multiple polyphenols and other ingredients. We have been studying the development of targeted delivery of drugs to the site of aneurysms so that a minimal dose of drug will be locally delivered in a sustained release manner to not only prevent aneurysm development but to regress developed aneurysms, which is clinically more relevant. Here, we successfully demonstrate that such targeted delivery of pentagalloyl glucose (PGG) restores degraded elastin, reduces MMP activity and infiltration of inflammatory cells, and regresses already developed aneurysms in elastase-induced AAA model.

## Methods

### Preparation of DIR loaded albumin nanoparticles for *in vivo* targeting studies

DiR (1, 1-dioctadecyl-3, 3, 3, 3-tetramethylindotricarbocyanine iodide) (PromoCell GmbH, Heidelberg, Germany) loaded BSA (Seracare, Milford, MA) nanoparticles were prepared by a similar method as described previously^3 6^. Briefly, bovine serum albumin, BSA (250 mg) was dissolved in DI water (4 mL) and then DiR dye (25 mg suspended in acetone) was added to BSA solution and stirred for one hour following the addition of glutaraldehyde (EM grade 70%, EMS, PA, USA) at a concentration of 42 μg/mg BSA. The mixture was added dropwise to 24 mL of ethanol under sonication (Omni Ruptor 400 Ultrasonic Homogenizer, Omni International Inc, Kennesaw, GA) on ice for 30 minutes.

### Preparation of Pentagalloyl glucose (PGG) loaded BSA nanoparticles

PGG-loaded nanoparticles were obtained by dissolving 250 mg of bovine serum albumin, BSA (Seracare, MA) in 4 mL of deionized (DI) water as stated previously. PGG (125 mg) was dissolved in 400 μl of dimethyl sulfoxide (DMSO) then the solution was added to BSA solution. Glutaraldehyde solution at a concentration of 12μg/mg of protein (BSA) was added while stirring. After an hour of continuous stirring, the mixture was added dropwise to 24 mL of ethanol under continuous probe sonication. The sonication was continued for an additional 30 mins. The nanoparticles were separated by centrifugation and washed.

### Conjugation of the anti-elastin antibody to nanoparticles

PGG or DiR loaded BSA NPs were PEGylated (NHS-PEG (2000) Maleimide) (Avanti Polar Lipids, Inc., Alabaster, AL) to achieve a sulfhydryl-reactive particle system. A polyclonal anti-elastin antibody (custom-made at Clemson University) was thiolated with Traut’s reagent (G-Biosciences, Saint Louis, MO) according to the manufacturer’s protocol. Thiolated antibodies obtained were added to the PEGylated NPs (4 μg antibody per 1 mg NPs) and incubated overnight.^7 3^ The PGG loaded and elastin antibody conjugated nanoparticles are named as EL-PGG-NPs, while DiR loaded NPs are named as EL-DiR-NPs.

To study NP targeting in vivo, we also made elastin antibody conjugated gold nanoparticles (EL-GNPs) as described previously^8^. (details are presented in the supplement).

### Elastase mediated AAA in mice

AAA was induced in specific pathogen-free C57BL/6 background male mice obtained from The Jackson Laboratory (Bar Harbor, ME). Mice were accommodated 3-5 per cage, at 22-24°C, 40-55% humidity and 12-hr light/dark-light cycle. The studies were carried out with approval from the Clemson University Institutional Animal Care and Use Committee (IACUC) following the guidelines of the Clemson University Animal Research Committee. Mice receive humane care in compliance with NIH Public Law 99-158.

Briefly, to induce AAA, the mouse was anesthetized with isoflurane, and a laparotomy exposed the abdominal aorta. Porcine pancreatic elastase (PPE; Sigma-Aldrich Co., St. Louis, MO, 7.6 mg/ml) was applied peri-adventitially for 12 mins, and subsequently, the aorta was rinsed with sterile PBS^9, 10^. After the abdominal specimens were placed in the original order, the fascial layers were closed with sutures. Each mouse was transferred to a new cage and was closely monitored for 24 hrs. Progression of disease in animals was monitored via high-frequency ultrasound imaging (Vevo2100, VisualSonics, Toronto, Canada). Sham group aortae were treated with PBS (no elastase).

After the surgery, mice were kept for two weeks to allow aneurysm development. Two weeks after elastase application, when significant aneurysms were developed, animals received two tail-vein injections of EL-PGG-NPs (10 mg/kg body weight) one week apart. Freshly prepared particles were delivered in sterile PBS. Control animals did not receive any therapy (n-10 per group). All animals were sacrificed at four weeks.

Details of animal study design are shown in Figure 1.

**Legend Figure 1.**
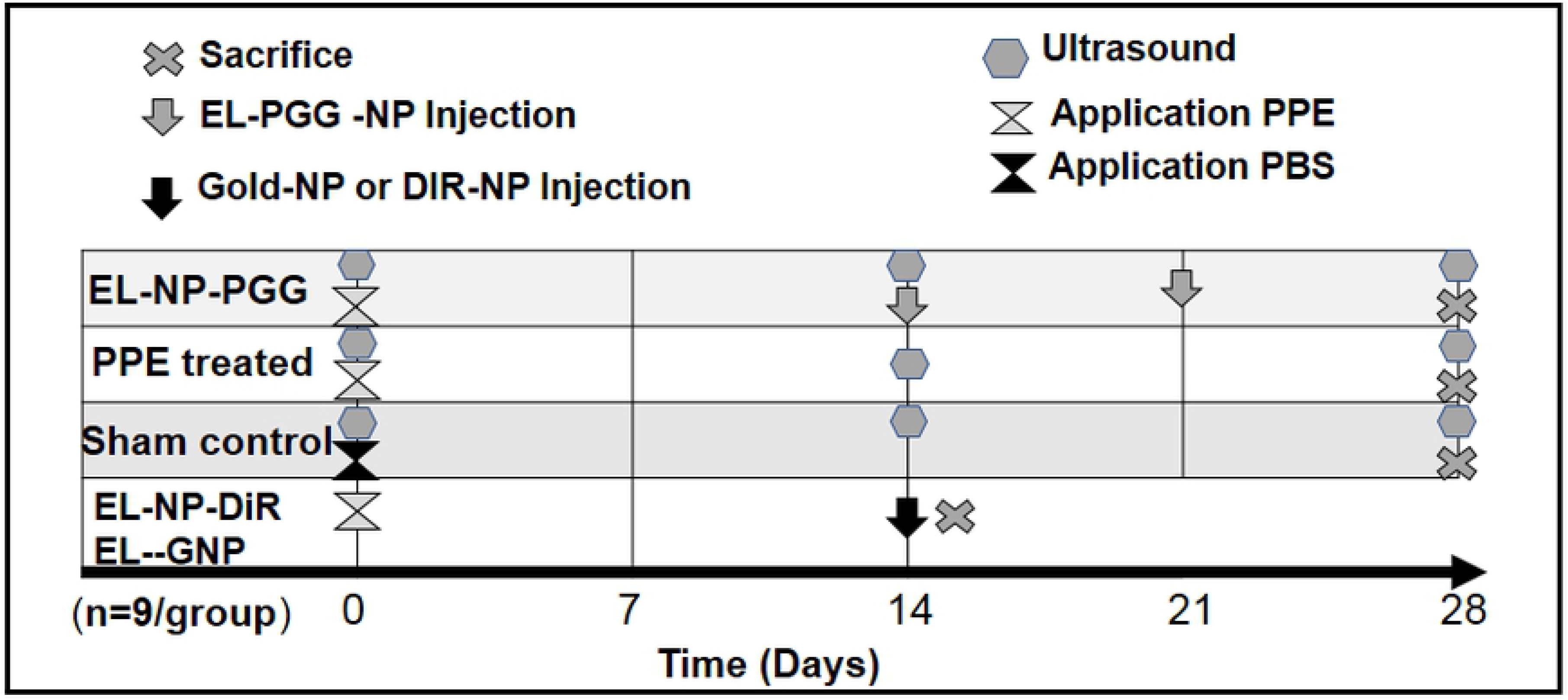
Schematic of study design

### Ultrasound analysis of the aneurysms

The percentage dilation and the circumferential strain of the aorta were assessed by using the ultrasound system. The animals were anaesthetized by carefully supplying 1% to 3% isoflurane during imaging and placed on the imaging table in a supine position. Mouse heartbeat rate and body temperature were also monitored during the imaging process. Sagittal and transverse images of aortas were obtained in motion mode, B mode and colour doppler mode. Systolic and diastolic inner diameters were measured and recorded at three different regions on each aneurysm or parent vessel using the built-in ultrasound software. The diastolic-to-systolic circumferential Green-LaGrange strains were analyzed using axial symmetry. The Circumferential Strain was calculated using the equation:

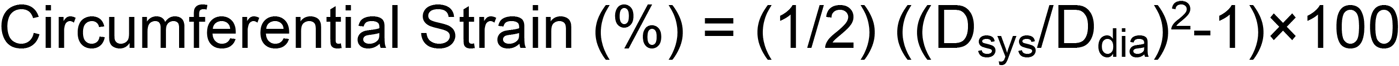

Where D_sys_ represents the inner systolic aortic diameter, and D_dia_ represents the inner diastolic aortic diameter.

Inner aortic diameters were measured on the abdominal aorta at the PPE applied region of the aorta by ultrasonography in basic-mode-images at three different time points during a cardiac cycle. A morphological aortic size assessment of explanted aorta was performed of the aneurysmal abdominal aortic region. Outer aortic diameters were visually measured after laparotomy. Mean values were then calculated for each aorta. The dilation was calculated using the equation given below, Percentage change in diameter of aorta was calculated using the equation as a percentage change (%) = ((D2/Dd1)/D1) × 100 Where, D1= initial aortic diameter, D2= final aortic diameter

### Histology of aorta sections

Formalin-fixed 2-3 mm aorta pieces were processed in Tissue-TEK^®^ tissue processor (Sakura Finetek USA, Inc., Torrance, CA) overnight and embedded in paraffin. Five-micrometer aorta sections were mounted on positively charged glass slides and were baked overnight to let the tissue attach to the slide and melt paraffin. Then the slides were deparaffinized with xylene following hydration with graded ethanol. Slides with aorta sections were stained for Verhoeff-van Gieson (VVG) (Richard-Allen Scientific, San Diego, CA) to visualize the elastin fibers. Overall tissue morphology was assessed by staining aorta sections with hematoxylin and eosin (H & E).

### Immunohistochemical analysis of aorta sections

Paraffin-embedded aorta sections mounted slides were subjected to heat-induced antigen epitope retrieval with citrate buffer (Thermo Scientific, MA). The slides were incubated overnight at 4°C with primary antibodies for anti -MMP-9 (Rabbit anti-Mouse) (Invitrogen, Carlsbad, CA), Rabbit anti-Mouse anti -MMP-2 (R & D Systems, Minneapolis, MN), Mouse anti-CD-68 (Novus, Cambridge, United Kingdom), anti-Mouse Mac-2 (Cedarlane, Burlington, Canada), and anti-mouse TGFβ1 (R & D Systems, Minneapolis, MN). The sections were incubated with relevant secondary antibodies (Goat anti-Mouse) (Invitrogen, Carlsbad, CA). IHC staining was completed with IHC kit (Enzo Life Sciences, NY). Slides were visualized by 3-Amino-9-ethylcarbazole (AEC) (Vector Laboratories, Burlingame, CA) chromogens followed by an appropriate counterstain.

### Flow cytometry

Mice were anaesthetized, PBS perfusion was performed from the right ventricle to drain the blood from the vascular system, and the aortas were harvested, cleaned and minced into 2-to 3-mm pieces. Aorta pieces were incubated in 1-mL digestion solution {containing 1 mg/ml porcine pancreatic elastase (Sigma-Aldrich); Collagenase D, 0.2 mg/ml, sigma; and DNAse I, 0.2 mg/ml, Roche; in DMEM media} for one hour. Excess PBS was added to stop the enzymatic reaction, and the tissue was minced and meshed on 70 μm filters. Immunofluorescence staining was performed as our previous methods^11^. Briefly, cells were incubated with Fc block solution (purified anti-mouse CD16/CD32, clone 2.4G2, BD Biosciences) for 15 min at room temperature to prevent non-specific binding. For CD68 cell surface markers, cells were incubated with the fluorescently conjugated antibody APC anti - CD68 (clone FA-11) (BioLegend, San Diego, CA) in the dark for 30 min at 4°C. For intracellular cytokine TGFβ1 staining, cells were permeabilized with fix/permeabilisation buffer (eBiosciences, San Diego, CA) for 15 mins before antibody PE anti - TGFβ1 (clone TW7-2089) (BioLegend, San Diego, CA) staining. After the extensive washing of cells in FACS buffer, the cytometric acquisition was performed on an LSR II CytoFlex (Becton Dickinson). Data analysis was performed using FlowJo (TreeStar, Ashland, OR) software.

### Statistical analysis

Statistical analysis was conducted using GraphPad Prism 8 (GraphPad Software, San Diego, CA). Comparisons were performed by one-way or two-way ANOVA as appropriate. A posthoc test for multiple comparisons was performed. The significance was determined as p < 0.05. Logarithmic transformation was conducted before statistical analysis for percentage change for statistical analysis.

## Results

### Confirmation of nanoparticle targeting to the AAA

To test if elastin antibody conjugated nanoparticles target AAA site, first, we confirmed the delivery of EL-GNPs (with micro-computed tomography, microCT) and EL-DiR-NPs (with IVIS imaging). AAA was allowed to develop for fourteen days after elastase treatment (Figure 2A-2). At day-14, EL-GNPs and EL-DiR-NPs were injected through the tail vein. Twenty-four hours after delivery, EL-GNPs were detected at the site of AAA with 3D reconstructed microCT image of the PPE treated aneurysmal aorta (Figure 2A-4) in the region of dilation of the aorta but no gold signal detected in sham control aorta (Figure 2A-3). Similar targeting of NPs to AAA site was found for EL-DiR-NP group with fluorescence located at the AAA site (Figure 2A-6). in comparison with the sham control (Figure 2A-5). Histology showed elastin damage area co-localized with the signal of DiR-NPs (data not shown) similar to our previous study^3^. We measured the biodistribution of EL-DiR-NPs in kidneys, spleen, liver, lungs and aorta (Figure 2B). The highest percentage of targeting was observed in the aorta (~ 80% per tissue weight basis), confirming antibody bound NPs were effectively delivered to the AAA site.

**Legend Figure 2.**
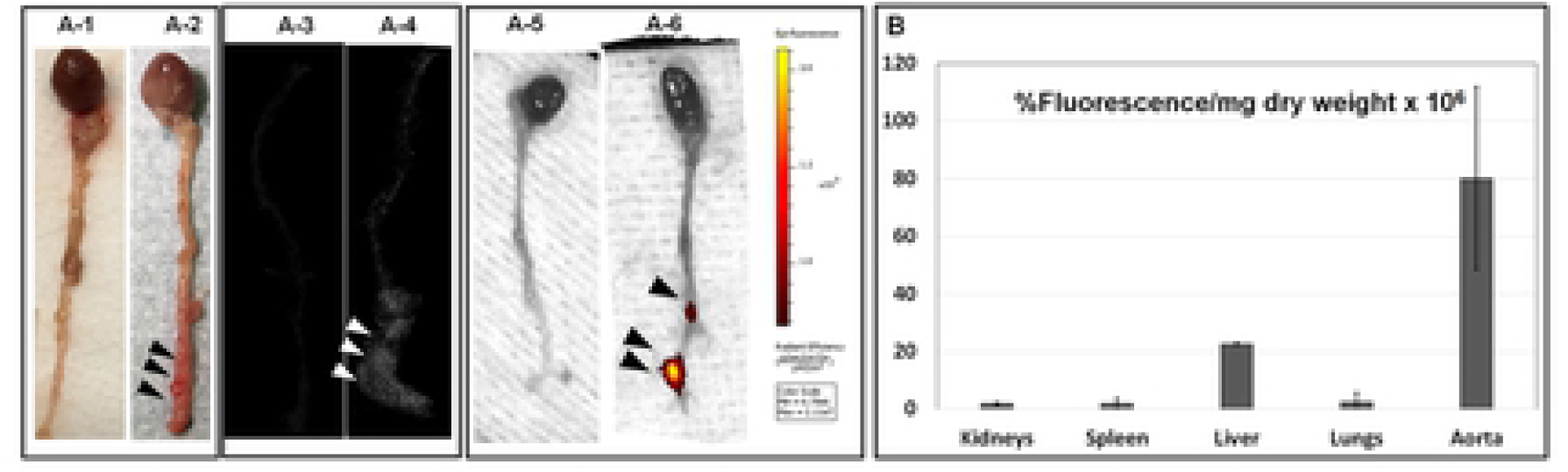
NP targeting to the aneurysmal aorta. **A-1 and A-2.** Representative picture of **the** explanted whole aorta from control and PPE treated mouse showing the aneurysmal region of the aorta. **A-3 and A-4.** Localisation of EL-GNPs within aneurysmal tissues at Day-14 with attenuation model of micro CT. The signal of EL-GNPs is visibly stronger in the aneurysmal aorta which is localized with the expanded region. The control aorta is devoid of signal. **A-5 and A-6.** IVIS images of control and PPE treated mouse aortas. The fluorescence for EL-DiR-NPs is visibly strong in PPE applied regions of aorta showing targeting to degraded elastin, while the PBS applied aorta is devoid of fluorescence. **B.** Bar graph showing biodistribution of EL-DiR-NPs quantified by the percentage of fluorescence per mg of dry tissue in different organs measured by IVIS imaging. The fluorescence was found significantly localised in the aneurysmal aorta confirming the targeting of nanoparticles.

### PGG delivery regresses already developed aneurysms

The internal aortic diameter of PPE treated mice significantly increased above 75% at day-14, and it continued to increase up to 150% at day-28 (Figure 3A). At day 14, when AAA was developed, systemic delivery of EL-PGG-NP (once a week for two weeks) lead to a significant decrease in the internal diameter (below 20%) (Figure 3B).

**Legend Figure 3.**
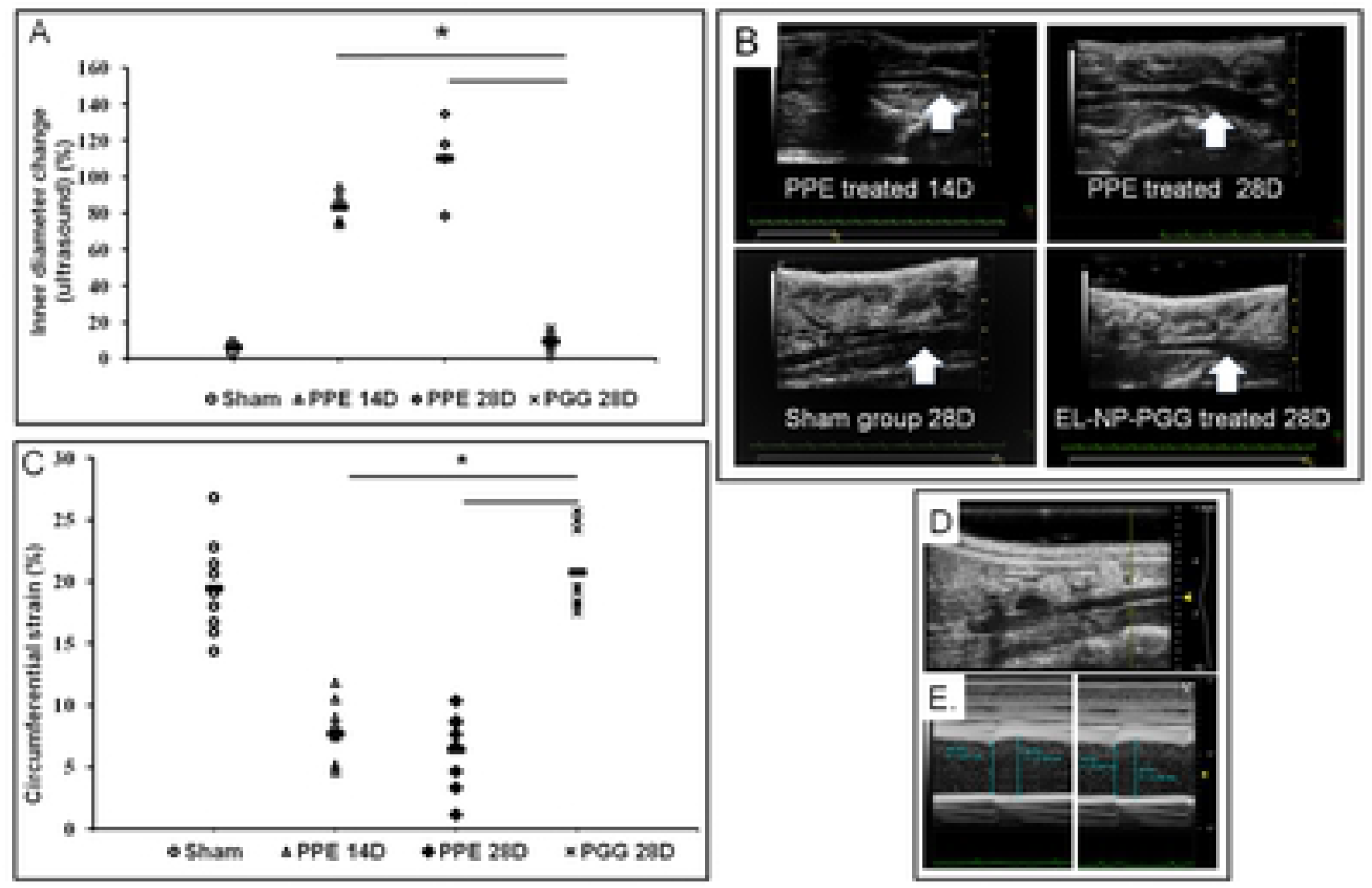
Reversal of aneurysm after PGG nanoparticle treatment. Representative data of internal diameter of the aorta as measured by ultrasound. **A.** Representative scatter plot showing the internal diameter of the aneurysmal region of abdominal aorta decreases significantly after the PGG nanoparticle treatment (EL-PGG-NP) in comparison to elastase treated control (PPE) group. * p < 0.0001 as compared to control. **B.** A representative B-mode ultrasound *in vivo* scan of aortas showing the internal diameter decreases in EL-PGG-NP group. The size of aneurysm increased from Day-14 to Day-28 in control group, which is noticeably decreased after PGG nanoparticle treatment. **C.** Scatter plot of circumferential Green-LaGrange strains throughout the cardiac cycle. In EL-PGG-NP group, the circumferential strain of aneurysmal aortas increased significantly in comparison to the PPE treated control aorta at Day-28 (p < 0.0001) showing improved elasticity. **D and E.** Representative M-mode *in vivo* ultrasound images of abdominal aorta showing the measurement of systolic and diastolic diameter.

The circumferential strain of the PPE treated aneurysmal aorta significantly decreased both at day 14^th^ and 28^th^ (Figure 3C) suggesting the loss of elasticity. After the intravenous delivery of EL-PGG-NP, the circumferential strain significantly increased in comparison to PPE treated on day 28 and reached to sham control values (Figure 3C). The representative ultrasound images show how the circumferential strain was obtained at diastole (Figures 3D and 3E).

The gross pictures of the aorta after laparotomy at day-28 explant showed increased external diameter and significant inflammation in the PPE group (Figure 4A). The outer diameter of the aneurysmal aortas increased significantly above ~ 95 % at day-14 and up to ~200 % at day-28 in the PPE treated mice (Figure 4B). The histology at 14-day explant confirmed aneurysmal expansion and significantly degraded elastic lamina in PPE treated group (Figure 4C). In EL-PGG-NPs group, the external diameter significantly decreased below ~ 45 % in comparison to PPE treated group at day-28 (Figure 4B). These data suggest that site-specific delivery of PGG with nanoparticles reverses the already developed aneurysm and restores arterial mechanics^12 13 14, 15^.

**Legend Figure 4.**
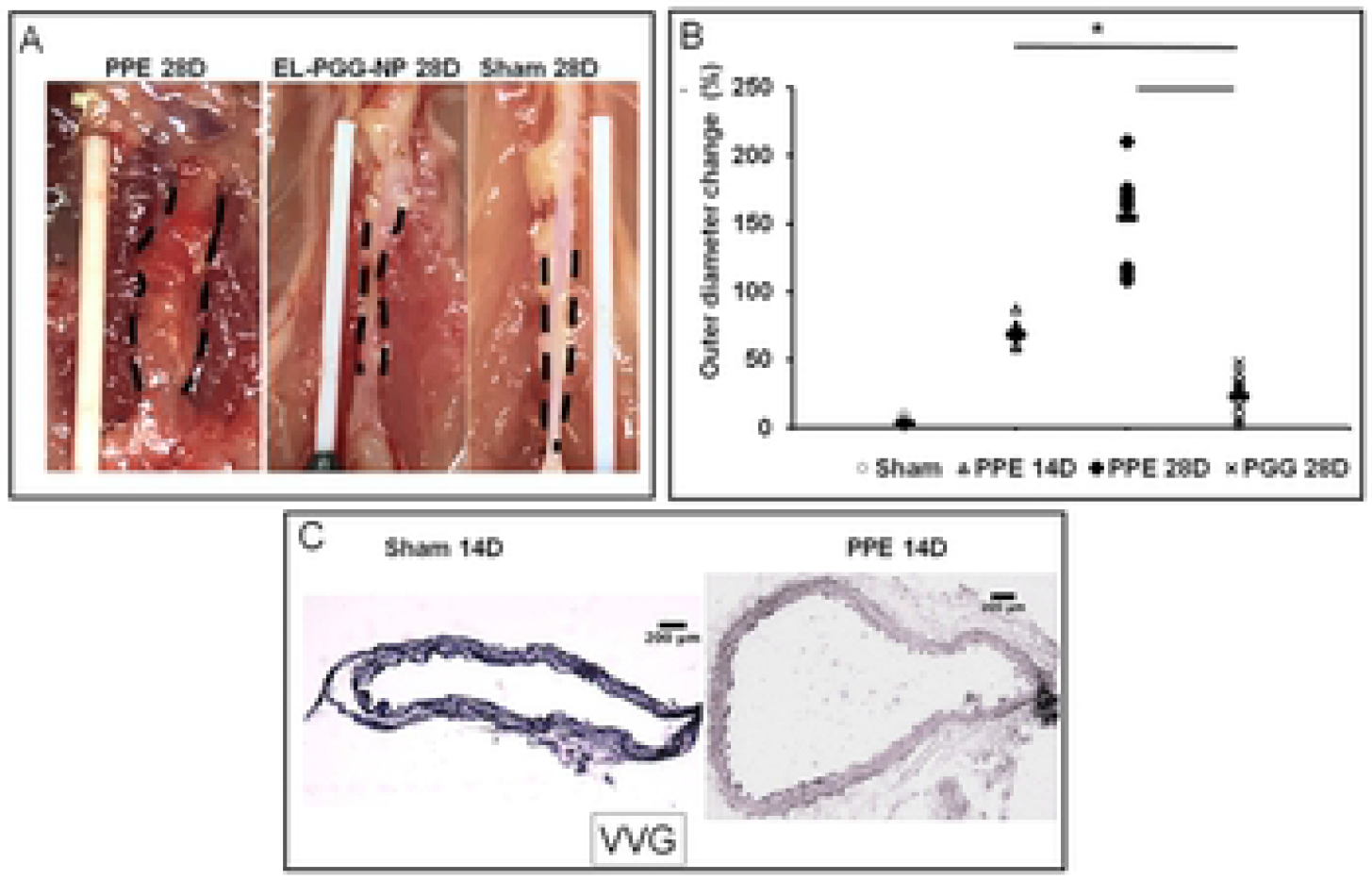
PGG nanoparticle treatment decreases the external diameter of aorta and adventitial inflammation. **A.** Representative gross pictures of aortas at 28 d. The aorta is visibly dilated in PPE treated mouse at the abdominal aortic regions with a significant inflammatory capsule. The PGG nanoparticle treated mouse aortae (EL-PGG-NP) were not aneurysmal and looks similar to sham controls (n=10 per group). **B.** The outer diameter of PPE treated aorta expanded up to 70% at 14d, and continue to expand to ~150% at day 28. After the two treatments of PGG nanoparticles starting at day 14, the external diameter of the aorta significantly decreased. * p < 0.0001 as compared to PPE treated controls. **C.** Representative histological sections (VVG stain for elastin-black) of PBS and PPE treated mouse aorta sections showing significant elastin damage at Day-14 at which point the PGG nanoparticle therapy was initiated.

### PGG delivery restores elastin and decreases inflammatory cell infiltration in the aorta

Histology with VVG stain showed significant degradation of elastin in the PPE treated aorta section compared with sham at 28-day explant (Figure 5A). In EL-PGG-NP group elastic fibers were restored in the media (Figure 5A). In H & E staining PPE treated aorta section was found thickened with infiltration of inflammatory cells in the aorta (Figure 5B). In EL-PGG-NP group, darkly pigmented nuclei of cells are sharply decreased implicating the decrease in the infiltration of inflammatory cells (Figure 5B).

**Legend Figure 5:**
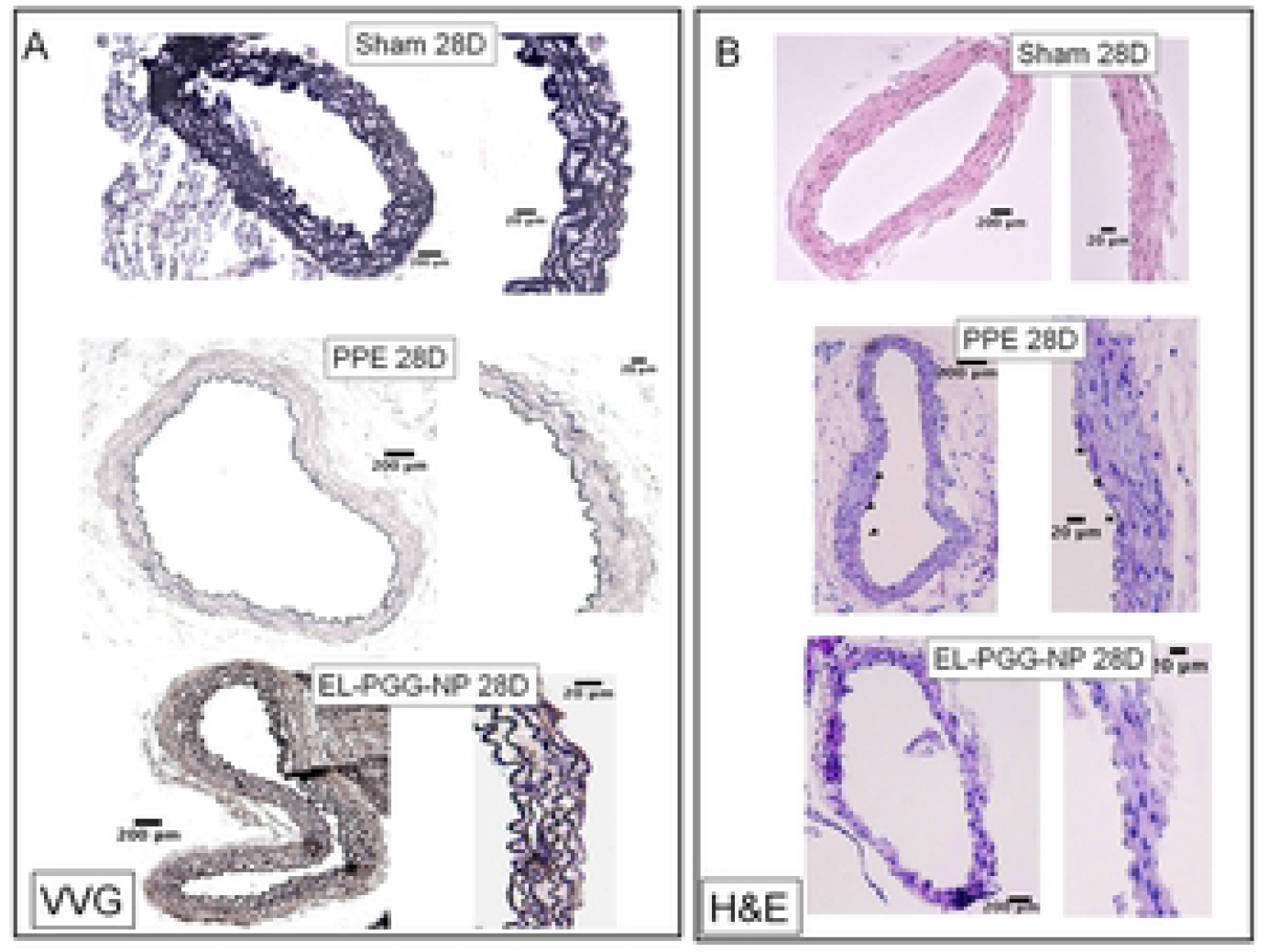
Restoration of elastic lamina after PGG nanoparticle therapy. **A.** VVG staining at 28d explants shows significant degradation of the elastic lamina in controls and regeneration of elastic lamina in the PGG treatment group (EL-PGG-NP). **B.** Hematoxylin and Eosin staining of PBS (Sham), PPE, and EL-PGG-NP group mouse aorta sections. Elastase treatment (PPE) caused significant adventitial inflammation while EL-PGG-NP group showed normal aorta similar to sham.

### PGG delivery decreases Matrix metalloproteinase (MMPs) in the aorta

MMPs degrade ECM, specifically elastin, and play a key role in the development of AAA in aorta^16 17^. These MMPs are secreted by inflammatory cells that are infiltrated in the aneurysmal aorta^18^. A significant amount of MMP-9 and MMP-2 on IHC staining were observed on PPE treated aorta sections both in adventitia and media of the aorta. Those levels were significantly decreased in the EL-PGG-NP treatment group (Figure 6A and 6B). These results indicate that the systemic delivery of EL-NP-PGG for two weeks decreases matrix metalloproteinases (MMPs) especially MMP-9 and MMP-2 activity.

**Legend Figure 6:**
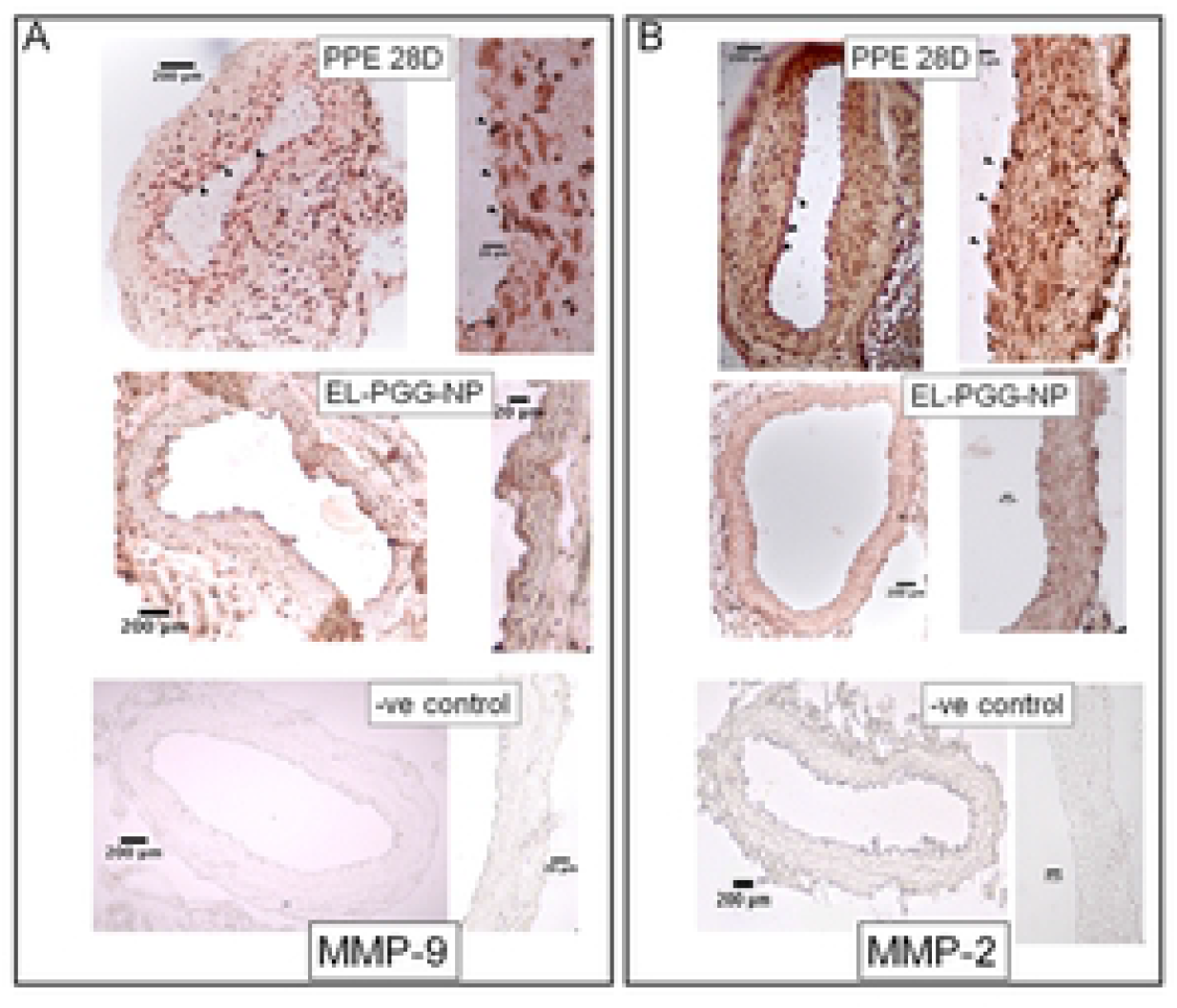
Reduction in MMP-2, MMP-9 and Mac-2 after the treatment of PGG nanoparticles. A. IHC staining of elastase treated (PPE), PGG nanoparticle treated (EL-PGG-NP), and PBS (Sham) treated mouse aorta sections, PGG treatment decreased MMP-9 staining in comparison to PPE treated aorta sections. **B.** PGG treatment also decreased MMP-2 staining in comparison to PPE treated mouse section. **C.** IHC staining for Mac-2 macrophages in mouse aorta sections showing PGG treatment decreases the infiltration of macrophages in the medial layer of the aorta in comparison to that in PPE treated aorta.

### PGG treatment decreases macrophage cells in the aorta

Immunofluorescence staining was conducted for CD 68 cells, which is also known as pan-macrophage and is regarded as M1 macrophage marker^19^. We observed that after the treatment with PGG, CD 68 positive cells significantly decreased in the aneurysmal aorta in comparison to PPE treated aorta (Figure 7A). Similarly, CD 68 positive cells were significantly decreased in aneurysmal aorta measured by immunofluorescence staining (Figure 7C). Moreover, a massive number of activated macrophages of Mac-2 positive cells observed in the PPE treated aortic sections, while the Mac-2 positive cells significantly decreased in EL-PGG-NP group (Figure 7B), especially in medial layer. These results suggest that the PGG treatment reduces the macrophage cells in the aorta that may be due to the decrease in overall inflammation in the aorta.

**Legend Figure 7:**
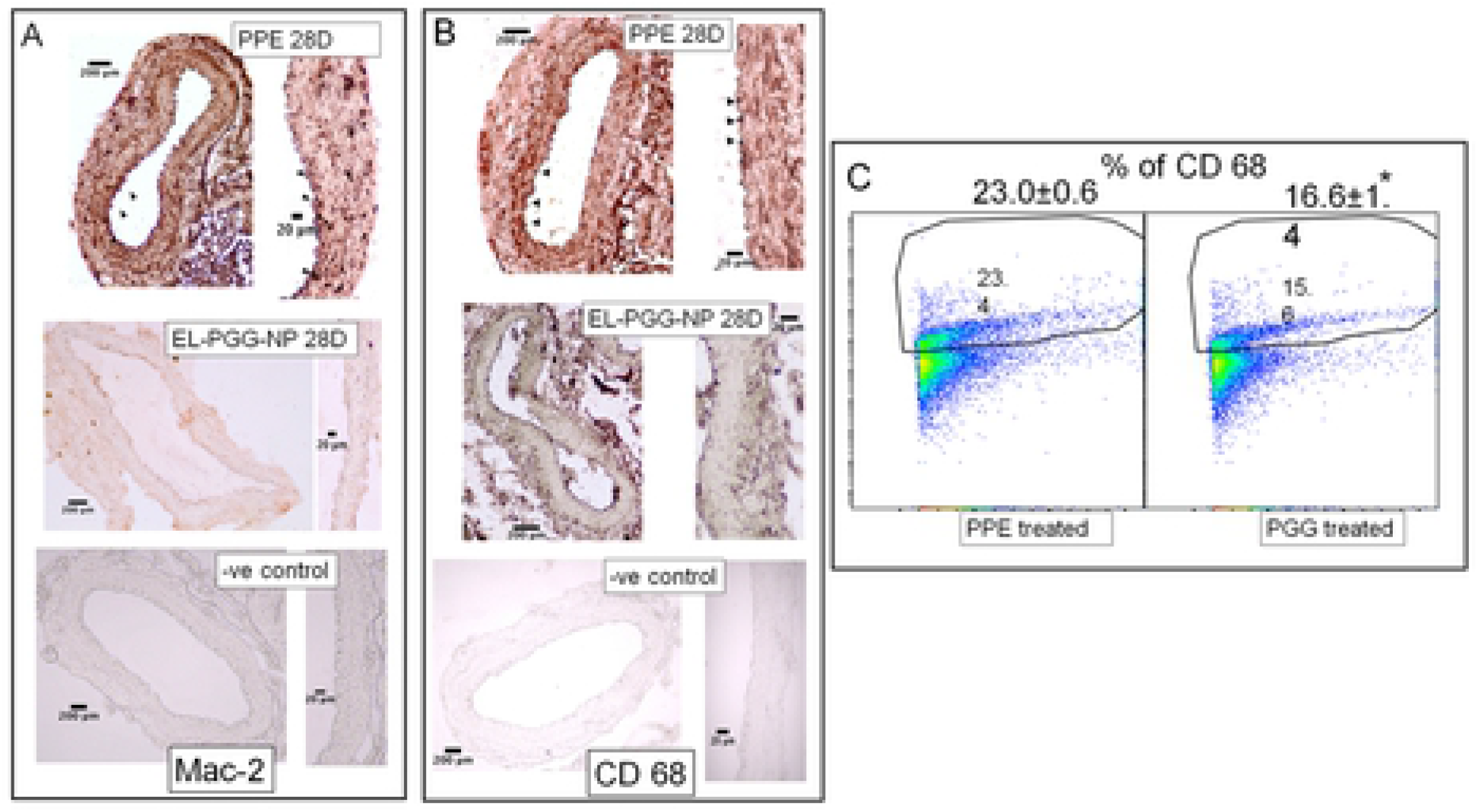
Reduction of CD 68 macrophages in the aorta of the PGG nanoparticle treatment. **A.** IHC staining of PPE, PGG (EL-PGG-NP), and PBS treated mouse aorta section showing PGG treatment decreases the infiltration of CD 68 macrophage in the medial layer of aorta section. **B.** Flow cytometry scatter plot showing PGG treatment significantly decreases CD 68 macrophage in PGG treated aorta region in comparison to that in PPE treated aorta. * p < 0.001.

### PGG treatment Decreases TGFβ1 in the aneurysmal aorta

TGFβ1 plays a key role in the progression of aneurysm. Others have shown that TGFβ1 administration exacerbates aneurysms^20 9^. IHC staining for TGFβ1 was performed to assess the presence of TGFβ1 in the aneurysmal aorta, in PPE and PGG treated aorta sections. We observed that TGFβ1 was heavily present in the PPE treated aneurysmal aorta (Figure 8A). We found that after the treatment with PGG nanoparticles (EL-PGG-NP group), flow cytometry showed significant decrease in intracellular TGFβ1 in the aneurysmal aorta in comparison to PPE treated aorta (Figure 8B).

**Figure 8:**
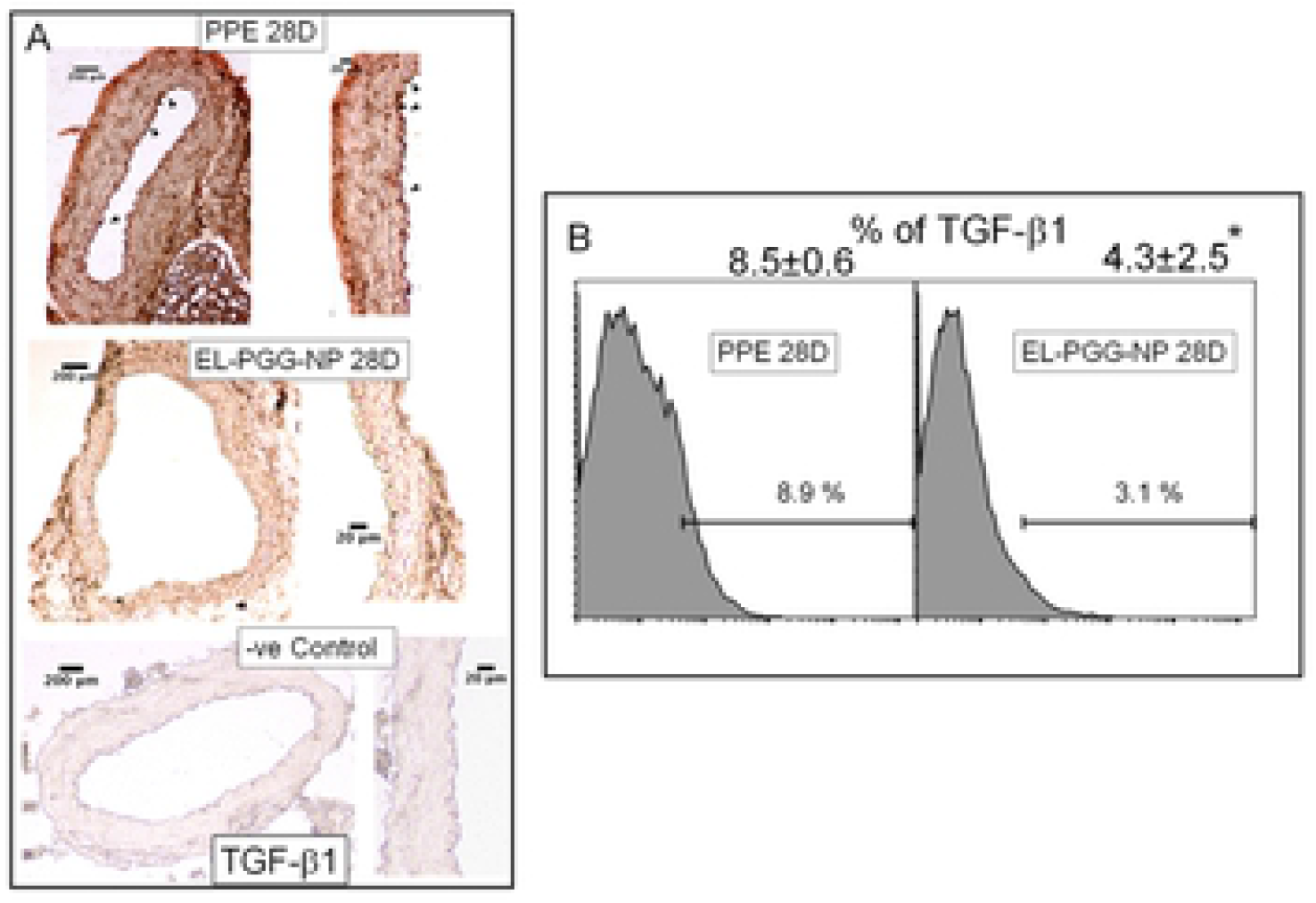
A reduction of TGFb1 after PGG nanoparticle treatment. A. IHC staining of aorta section showing PGG treatment drastically decreased the staining of TGFb1 in aorta section in comparison to PPE treated aorta. **B.** Flow cytometry histogram showing PGG treatment significantly decreases the percentage of TGFb1 in comparison to that in PPE treated aneurysmal aorta. * p < 0.0001as compared to PPE treated.

## Discussion

The purpose of this study is to test if our novel targeted NP based PGG delivery that only targets degraded elastin could reverse already developed aneurysms in elastase-induced AAA. We not only show that such targeted delivery regresses aneurysms, but we also show that such therapy restores elastin lamina, decreases matrix metalloproteinase (MMPs), immunomodulating cytokine TGFβ1, and infiltration of activated macrophages at the aneurysmal site.

Elastin is found in the major blood vessels of nearly all vertebrates formed of a pulsatile, high-pressure closed circulatory system^21^. Elastic fiber imparts reversible distensibility to the large arteries, allowing the aorta to deform during cyclic hemodynamic loading, with no permanent deformation or energy dissipation upon load retrieval^22^. One of the significant consequences of aneurysm development is degradation elastic lamina. Unfortunately, adults cells cannot regenerate lost elastic fibers on their own as the microfibrillar assembly is lacking^23^. Furthermore, degraded elastic lamina can release elastin degradation products (EDPs) that are chemokines for inflammatory cells and modulate M1/M2 macrophage polarization^24^. Thus, stopping elastin degradation is a crucial step in preventing AAA development. We previously reported local PGG treatment of stabilized arterial elastin and prevented degradation of elastin^12^. We have also shown that PGG treatment of vascular smooth muscle cells allowed the development of crosslinked elastin in cell cultures^4^. In the calcium chloride rat AAA model, dual-targeted therapy reversed calcification and aneurysmal expansion^3^.

Here we chose a more aggressive model of AAA where the local application of elastase significantly degraded elastin in the medial layers by day-14. We applied porcine pancreatic elastase (PPE) for 12 mins, which developed variable dilation after two weeks but showed significant elastic lamina degradation in all mice. Bhamidipati et al. used PPE for 10 mins, and with an oral dose of doxycycline showed prevention of the progression of AAA^25^. Similarly, Wang et al. applied PPE for 30 mins and used an oral dose of grape seed polyphenol to show prevention of AAA development. However, the amount of grape seed polyphenol given daily (400-800 mg/kg) is excessive (correspond to 28-55 gm dose per day in humans)^10^. Similarly, Setozaki et al. showed green tea polyphenol Epigallocatechin-3-gallat (ECGC) given in drinking water prevented AAA formation and reduced elastin degradation and inflammation in a combination of elastase and CaCl_2_ mediated AAA in rats. They also used excessive amounts (correspond to 60-80L green tea consumption as it has low ECGC). Furthermore, most previous studies initiated therapies at the onset of elastase treatment to prevent AAA formation and did not target drugs to the site of the aneurysm. We wanted to find out if targeted therapy could reverse already developed AAA in this aggressive model. Such targeted treatment would need a small amount of drug released at the site to reverse aneurysms.

We used our novel nanoparticles coated with elastin antibody that only targets degraded elastin while sparing healthy elastin to deliver PGG to the AAA site^3, 6, 13, 26^. We chose albumin-based NPs as they are proven to be non-toxic^27^ and are used to deliver paclitaxel in patients (Abraxane). We previously demonstrated that EL-PGG-NPs have no hepatic toxicity for systemic delivery and these particles are targeted at the aneurysmal site^3^.

Here we clearly show that nanoparticle reach and adhere to the AAA site in the elastase model. Both albumin-based and gold nanoparticles could be targeted at the AAA site. Next, we delivered PGG loaded nanoparticles on day-14 to test if PGG delivery could restore elastin that is already degraded by elastase treatment. We performed two systemic injections of nanoparticles one week apart as we have shown that PGG slowly releases from particles over a period of several weeks, and nanoparticles stay at the site for more than a week^13^. Histology after two weeks of PGG delivery showed elastic laminae in the medial layers were restored in the aneurysmal aorta in all PGG treated mice (n=10), while it was severely degraded at the onset of our therapy at day-14. Such restoration of elastic lamina also led to a significant reduction in the aneurysm size shown by both internal and external aortic diameter measurements. We also demonstrated that PGG significantly increased aortic circumferential strain. The previous report shows aneurysmal aorta’s circumferential strain is inversely correlated with the elastin fiber loss and directly correlated with the diameter decrease^28^. Thus, not only PGG treatment restored elastic lamina and reversed AAA but allowed aorta to restore properties to that of the healthy aorta.

During the progression of AAA, accumulation, and activation of inflammatory cells takes place in the aorta. These inflammatory cells play a key and substantial role in vascular remodeling^9, 29^. In our study, we found a significant amount of CD 68 (M1 type macrophage) and activated macrophage Mac-2 in PPE treated aorta. Macrophages rapidly secrete a wide range of inflammatory mediators such as IL-13, IL-6, and TNFα and overexpression of these inflammatory mediators is reported in experimental AAA followed by the infiltration of monocytes, neutrophils and lymphocytes and then followed by VSMCs apoptosis^19,4^. Others have shown that apoptosis in VSMCs releases MMPs, which degrade ECM leading to vessel damage^30^. Macrophages are often found near collagen-producing cells, are key regulators of inflammation. They produce profibrotic mediators such as TGFβ1 and control the homeostasis of various matrix metalloproteinases and tissue inhibitors of matrix metalloproteinases^31 32^. In our study, after the treatment of PGG nanoparticles, CD 68 and activated macrophages Mac-2 drastically decreased in the aorta. Reduction of macrophages might have contributed to the reduction in inflammation and, thus, for the amelioration of aneurysm. Moreover, targeted PGG therapy decreased the local proliferation CD 68 positive macrophage cells in the already formed aneurysms. Thus, not only PGG delivery restored vascular elastin, but it also reduced inflammatory cytokines and macrophages at the site of AAA. We hypothesize that PGG binding to elastin stops further degradation of elastin and allows tropoelastin molecules that are secreted by VSMCs to coacervate and create new elastic fibers. We showed it before that PGG binds strongly to elastin and prevents its further degradation^12^. Such binding can avoid generation of elastin peptides that are known chemoattractant to macrophages^33^.

In our study, we observed MMP-9 and MMP-2 decreased after the treatment of PGG nanoparticles. MMP-9 and MMP-2 are required for the development of AAA since MMP-9 and MMP-2 knockout mice are resistant to AAA disease^34^. Others have shown that polyphenols like curcumin and xanthohumol inhibit matrix metalloproteinase activity and even have anti-inflammatory effects^35, 36^. Elastase treatment of aorta leads to the recruitment of inflammatory cells. Recruitment of macrophages and monocytes in aorta regulates various matrix metalloproteinases. These inflammatory cells secrete MMPs in the aorta during AAA^17^. The MMP activity reduction with PGG could have occurred in two ways. We have previously shown that PGG inhibits MMP activity in vitro by gel zymography^37^. It is also possible that the prevention of macrophage recruitment is due to inhibition of elastin fragmentation by bound PGG to elastin, as shown above, which can decrease local MMP activity.

Immunomodulatory cytokine TGFβ1 plays a key role in ECM modulation. TGFβ1 neutralization either worsened or mitigated aneurysm formation, depending on whether treatment was initiated before or after the establishment of the aneurysm in a thoracic aorta^20^. TGFβ1 contributed to the progression of the aneurysm when administered after the formation of the aneurysm in the experimental mouse model^20 9 38^. Depending on circumstances, TGFβ1 signaling may follow the Smad2 dependent or independent pathway for its effect on the pathogenesis of aneurysm formation. Healthy elastic fibers are known to sequester TGFβ1 through latent TGFβ binding protein (LTPBs) that is associated with microfibrils of elastic fiber^39^. Thus, healthy extracellular matrix modulates TGFβ activity, and loss of LTPBs due to degradation of elastin can increase the activity of TGFβ1. In this research, we observed that significant levels of TGFβ1 and Mac 2 were present in the PPE treated aneurysmal aorta. We found that after the treatment of PGG nanoparticles, TGF β1 was significantly decreased in the aorta. That might be due to the reduction of monocyte and macrophage infiltration in the aorta. Research has shown that monocytes and M2 phenotype macrophages secrete TGF β1^40 31^. It is possible that the restoration of the healthy matrix in the aorta led to a decrease in TGFβ1 activity in the aorta. These data suggest that PGG may further decrease the pathogenicity in aneurysm by decreasing TGFβ1 signaling in AAA^9 38^. It is noteworthy that a minimal amount of PGG was required for the reversal of AAA. We used 10 mg/kg nanoparticle dose, and PGG loading in nanoparticle was 30%. Thus, only 3 mg PGG was used two times in two weeks with effective dose of ~425 μg/day/kg for two weeks; thus, a small locally targeted PGG can be effective in reversing AAA.

In conclusion, in our study, site-specific delivery of PGG with our targeted nanoparticles that accumulate in the AAA site reversed already formed aneurysms, regenerated elastic lamina, reduced macrophages, and inflammatory cytokines and MMPs in the aorta. The therapy also restored the healthy mechanics of the aorta. Such therapy can be translated to patients who already have AAA disease as there are no therapies available to reverse AAA.

## Acknowledgments

We appreciate the help of staff of Godley Snell Research Center animal facility of Clemson University for helping in animal study. We thankfully acknowledge Dr. James Morris lab for letting us to use flow cytometer supported by NIH grant P20GM109094.

## Abbreviations

AAA: Abdominal aortic aneurysm
BSA: Bovine serum albumin
DIR: 1,1-Dioctadecyl-3,3,3,3-tetramethylindotricarbocyanine iodide
EL-GNP: elastin antibody conjugated gold nanoparticle
EL-NP: Elastin antibody-conjugated blank nanoparticles
EL-PGG-NP: Elastin antibody-conjugated and PGG-loaded albumin NPs
FACS: Fluorescence-activated cell sorting
MMP: Matrix metalloproteinase
NPs: Nanoparticles
PGG: Pentagalloyl glucose
TGF-□: Tumor Growth Factor beta
IL: Interleukine

